# A torpor-like state (TLS) in mice slows blood epigenetic aging and prolongs healthspan

**DOI:** 10.1101/2024.03.20.585828

**Authors:** Lorna Jayne, Aurora Lavin-Peter, Julian Roessler, Alexander Tyshkovskiy, Mateusz Antoszewski, Erika Ren, Aleksandar Markovski, Senmiao Sun, Hanqi Yao, Vijay G. Sankaran, Vadim N. Gladyshev, Robert T. Brooke, Steve Horvath, Eric C. Griffith, Sinisa Hrvatin

**Affiliations:** Whitehead Institute for Biomedical Research and Department of Biology, Massachusetts Institute of Technology, 455 Main Street, Cambridge, MA 02142, USA; Department of Neurobiology, Harvard Medical School, Boston, MA 02115; Department of Neurobiology, Stanford University Medical Center, Stanford, CA, USA; Division of Genetics, Department of Medicine, Brigham and Women’s Hospital, Harvard Medical School, Boston, MA, 02115, USA; Division of Hematology/Oncology, Boston Children’s Hospital, Harvard Medical School, Boston, MA, USA; Department of Pediatric Oncology, Dana-Farber Cancer Institute, Harvard Medical School, Boston, MA, USA; Broad Institute of MIT and Harvard, Cambridge, MA, USA; Program in Neuroscience, Harvard Medical School, Boston, MA, USA; Epigenetic Clock Development Foundation, Torrance, CA, USA; Altos Labs, Cambridge, UK

## Abstract

Torpor and hibernation are extreme physiological adaptations of homeotherms associated with pro-longevity effects. Yet the underlying mechanisms of how torpor affects aging, and whether hypothermic and hypometabolic states can be induced to slow aging and increase health span, remain unknown. We demonstrate that the activity of a spatially defined neuronal population in the avMLPA, which has previously been identified as a torpor-regulating brain region, is sufficient to induce a torpor like state (TLS) in mice. Prolonged induction of TLS slows epigenetic aging across multiple tissues and improves health span. We isolate the effects of decreased metabolic rate, long-term caloric restriction, and decreased core body temperature (T_b_) on blood epigenetic aging and find that the pro-longevity effect of torpor-like states is mediated by decreased T_b_. Taken together, our findings provide novel mechanistic insight into the pro-longevity effects of torpor and hibernation and support the growing body of evidence that T_b_ is an important mediator of aging processes.

## Main

In response to food deprivation or harsh environmental conditions, many mammalian species engage energy conserving strategies, such as torpor and hibernation. Torpor is a state of profoundly decreased core body temperature (T_b_) and metabolic rate lasting from hours to days, whereas hibernation is a seasonal behavior comprising multiple bouts of torpor interrupted by periodic arousals to euthermia. These extraordinary adaptations raise many unanswered fundamental questions of homeotherm biology, one of the most compelling being the link between torpor and longevity. Natural torpor is characterized by tightly coupled, extreme physiological changes that have been individually implicated in aging and longevity, such as decreased core body temperature and metabolic rate, and caloric restriction. Indeed, hibernating species exhibiting long torpor bouts show extended longevity compared to closely-related non-hibernators, and longer lifespan than would be expected based on body-mass alone^1^. However, the lack of genetic tools and controllable systems to study torpor have left the central questions of how torpor affects aging, and whether hypothermic and hypometabolic states can be induced to slow aging and increase health span, largely unexplored.

The brain is known to control core body temperature and metabolism^2,3^. Specialized populations of neurons in the preoptic area (POA) of the hypothalamus receive information about physiological states, including peripheral and core body temperature, and modulate thermoregulatory circuits to maintain homeostasis^4–7^. In addition to the role of the POA in normothermic thermoregulation, recent work has identified neuronal populations in the POA whose activity is necessary for the normal expression of fasting-induced daily torpor in mice^8,9^. Importantly, stimulation of these neurons is sufficient to induce torpor-like hypothermic and hypometabolic states^8–12^. Building on this work, we develop a controllable torpor-like state (TLS) that can be safely maintained for days, and repeatedly induced over months, in non-transgenic laboratory mice. While torpor and hibernation are complex behaviors that TLS does not fully recapitulate, induction of TLS causes a profound decrease in core body temperature (T_b_), body temperature set point (T_set_), metabolic rate (VO_2_), respiratory quotient (RQ), activity, and food intake, phenocopying several key features of natural torpor and hibernation^13,14^.

To examine the effects of TLS on aging, we induce prolonged TLS to model a natural hibernation-like pattern over the course of months and find that TLS improved clinical measures of age-associated frailty in mice and slowed epigenetic aging in a tissue-specific manner. TLS had the greatest effect on epigenetic aging in the blood, where it slowed epigenetic aging by up to 76% in individual mice. Moreover, we leverage the uniquely controllable nature of our model to elucidate the underlying mechanisms of the pro-longevity effect of TLS on blood epigenetic aging, demonstrating that this effect is not mediated by caloric restriction nor decreased metabolic rate, but rather stems from a decrease in T_b_.

### Development of an inducible torpor-like state (TLS) in non-transgenic laboratory mice

Building on recent work that has identified the POA of the hypothalamus as a torpor-regulating brain region^8–12^, we tested whether targeted stimulation of the anterior and ventral portions of the medial and lateral preoptic area (avMLPA) of the hypothalamus could recapitulate key physiological features of torpor in non-transgenic laboratory mice.

Mice were implanted with telemetric temperature probes and stereotactically injected in the avMLPA with AAV-hSyn-hM3D(Gq)-mCherry, an adeno-associated virus (AAV) expressing a chemically activated receptor, Gq-DREADD (Gq-coupled Designer Receptor Exclusively Activated by Designer Drug) driven by the neuronally restricted *hSyn* promoter (Fig. 1a-f). Activation of neurons within the avMLPA via intraperitoneal injection of the Gq-DREADD-activating synthetic ligand clozapine-*N*-oxide (CNO) drove a decrease in T_b_ (Fig. 1a-f), consistent with targeting the avMLPA neuronal population whose activity is sufficient to induce a large decrease in T_b_ as observed during natural fasting-induced torpor bouts.

**Fig. 1.**
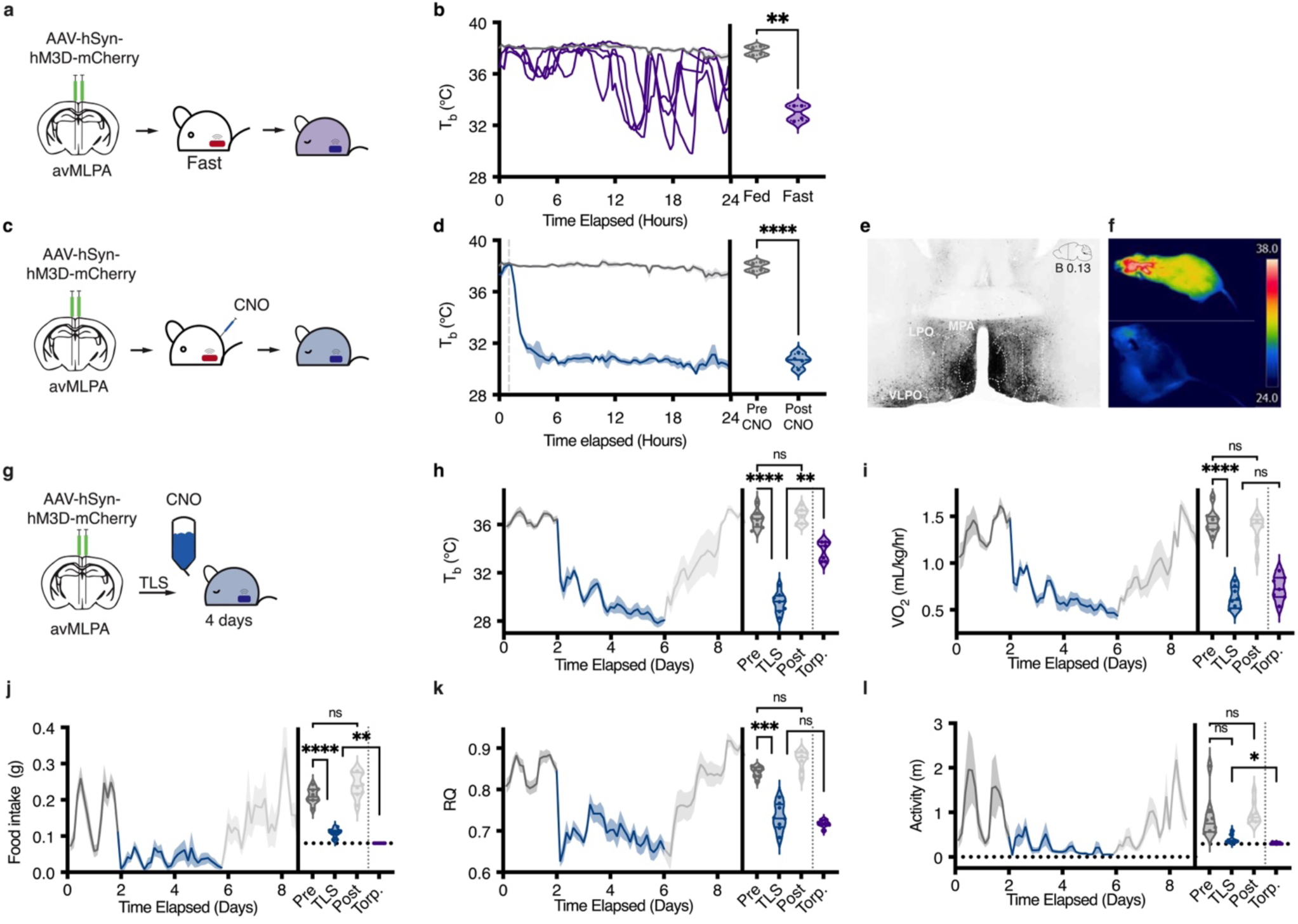
Development of an inducible torpor-like state (TLS) in non-transgenic laboratory mice. **a,** Schematic of injection of AAV-hSyn-hM3D-mCherry and subsequent fast. **b,** T_b_ over 24-hour fast (purple) as compared to baseline (grey). Baseline data plotted as mean ± SEM, fasting data plotted as individual mice. When fasted, all mice engaged in natural fasting-induced daily torpor as defined by T_b_ < 35 °C and not arousing. Mice began engaging in torpor bouts ∼10 hours into the fasting interval. During torpor bouts, individual mice had an average T_b_ (33.0 ± 0.32 °C) lower than at baseline (37.9 ± 0.18 °C) as determined by paired T-test (\*\**P* = 0.002) and reported as mean ± SEM. **c,** Schematic of injection of AAV-hSyn-hM3D-mCherry and subsequent CNO injection. **d,** T_b_ at baseline (grey) and after stimulation with CNO (blue) (*n* = 4), CNO injection indicated by dashed grey line. Data plotted and reported as mean ± SEM. Average T_b_ of mice 6-24 hours post-CNO injection was lower (30.6 ± 0.27 °C) than at baseline (37.9 ± 0.18 °C) as determined by paired T-test (\*\*\*\**P* < 0.0001). **e**, Representative coronal section displaying hM3D expression in the avMLPA as visualized by mCherry after viral injection. **f**, Thermal imaging of body temperature of a mouse at before (top) and after induction of TLS (bottom). **g,** Schematic of TLS induction through continuous administration of CNO via drinking water. **h-l,** T_b_, VO_2_, food intake, RQ, and activity at baseline (dark grey), during TLS (blue), and in recovery (light gray). Lines indicate mean, shading denotes ± SEM (*n* = 8). Violin plots graphed as the average of each individual at baseline, during TLS, in recovery, and during natural fasting-induced torpor bouts (purple, **Extended Data Fig. 1a-c**). All measured metabolic parameters had significant differences across conditions as quantified by one-way ANOVA adjusted for multiple comparisons by TUukey HSD (T_b_ Pre = 36.44 ± 0.26, TLS = 29.54 ± 0.30, Post = 36.71 ± 0.22, Torp. = 33.86 ± 0.30, \*\*\*\**P* < 0.0001) (VO_2_ Pre = 1.44± 0.04, TLS = 0.63 ± 0.04, Post. = 1.37 ± 0.05, Torp. = 0.73 ± 0.05, \*\*\*\**P* < 0.0001) (Food intake Pre = 0.16 ± 0.01, TLS = 0.03 ± 0.01, Post = 0.20 ± 0.02, Torp. = 0 ± 0, \*\*\*\**P* < 0.0001) (RQ Pre = 0.85 ± 0.01, TLS = 0.69 ± 0.02, Post = 0.89 ± 0.01, Torp. = 0.67 ± 0.0, \*\*\*\**P* < 0.0001) (Activity Pre = 1.11 ± 0.33, TLS = 0.20 ± 0.06, Post = 1.27 ± 0.17, Torp. = 0.03 ± 0.02, \*\**P* = 0.0015). Data reported as mean ± SEM.

In mice, fasting-induced daily torpor bouts last only several hours before arousal to euthermy^8,13,15–17^. Hibernators, by contrast, engage in torpor bouts lasting days to weeks^18–20^. Having found that targeted chemogenetic stimulation of the avMLPA recapitulates decreases in T_b_ as in fasting-induced daily torpor, we tested if this approach could be used to induce a torpor-like state (TLS) in mice lasting days to weeks, as seen in natural hibernation. Mice implanted with telemetric temperature probes and stereotactically injected in the avMLPA with AAV-hSyn-hM3D(Gq)-mCherry were continuously administered CNO via drinking water, inducing a hypothermic state that remarkably could be safely maintained for days at a time (Fig. 1g, h). To characterize and capture metabolic changes during TLS, we profiled T_b_, activity, food intake, metabolic rate (as measured by oxygen consumption (VO_2_)), and the respiratory quotient (RQ) using the Prometheon Metabolic System (Fig. 1h-l). We found that TLS caused a profound reduction across all measured parameters (\*\*\*\**P* < 0.0001), similar to changes seen in fasting-induced daily torpor (Fig. 1h-l, Extended Data Fig. 1a-c). Notably, over four days of TLS, the average body temperature dropped by nearly 7 °C, while metabolic rate and food intake dropped by ∼56% and ∼81%, respectively (Fig1. f-h). Upon removal of CNO from drinking water, mice spontaneously recovered across all measured metabolic parameters (Fig. 1h-l).

Natural hibernators achieve a lowered T_b_ during torpor by modifying key features of their thermoregulatory system, reducing both their sensitivity to decreased body temperature (*H*) and their theoretical set point temperature (*T_set_*) ^18–20^. To assess changes in these parameters in TLS mice, we monitored metabolic rate (VO_2_) and body temperature (*T_b_*) in these animals across varied ambient temperatures (T_a_) (8, 16, and 24 °C) (Extended Data Fig. 2a-e)^15^. Consistent with the entry into torpor of natural hibernators, we found that both *H* and *T_set_* were significantly reduced (Extended Data Fig. 2a-e) in TLS. Taken together, these results suggest that the thermoregulatory system during TLS shares key features with natural hibernation and fasting-induced daily torpor.

### TLS slows epigenetic aging across multiple tissues

Given the links between hibernation, which is comprised of multiple bouts of torpor interrupted by periodic arousals to euthermia, and longevity, we tested whether TLS slows aging. Aging is characterized by progressive decline in physiological function, increased susceptibility to disease, and death. While aging is a complex phenomenon, epigenetic clocks have recently emerged as powerful molecular biomarkers of aging^21–24^. Importantly, recent work across multiple species of natural hibernators has found that blood epigenetic aging is slowed during hibernation, suggesting that epigenetic clocks can capture torpor-related changes in aging processes^25,26^. We induced TLS to model a natural hibernation-like pattern by repeatedly administering CNO through drinking water, resulting in bouts of TLS with periodic arousals to euthermia (Fig. 2a, b). While on CNO, TLS mice had significantly lower T_b_ (30.96 ± 0.19 °C) than control mice (35.58 ± 0.02 °C) (\*\*\*\**P*<0.0001) (Fig. 2c, d). After three months, we analyzed the epigenetic age of the blood, liver, kidney, and cortex and found that TLS mice aged on average 80% less in the blood (\**P* = 0.007), and 20% less in the liver (\**P =* 0.034) than age-matched controls (Fig. 2e, f). In a secondary analysis, we applied additional epigenetic clocks, including two universal pan mammalian clocks which track highly conserved epigenetic aging effects. Again, we found that TLS mice had lower blood epigenetic age than control mice, evidencing the robustness of the effects of TLS on blood epigenetic aging (Extended Data Fig. 3a, b, c). There was no significant decrease in the epigenetic age of the kidney *(P =* 0.35) or cortex (*P* = 0.74) in TLS mice compared to age-matched controls (Fig. 2g, h) as corroborated by additional epigenetic and transcriptomic clock analyses (Extended Data Fig. 3a-d). Our results suggest that inducing TLS in a non-natural hibernator can recapitulate the effects of natural hibernation on blood epigenetic aging^26^. While work in natural hibernators has only explored epigenetic aging of the blood^26^ and skin^25^, both highly replicating tissues, our work across multiple tissues suggests that hypothermic and hypometabolic states may exert highly tissue-specific effects on epigenetic aging.

**Fig. 2.**
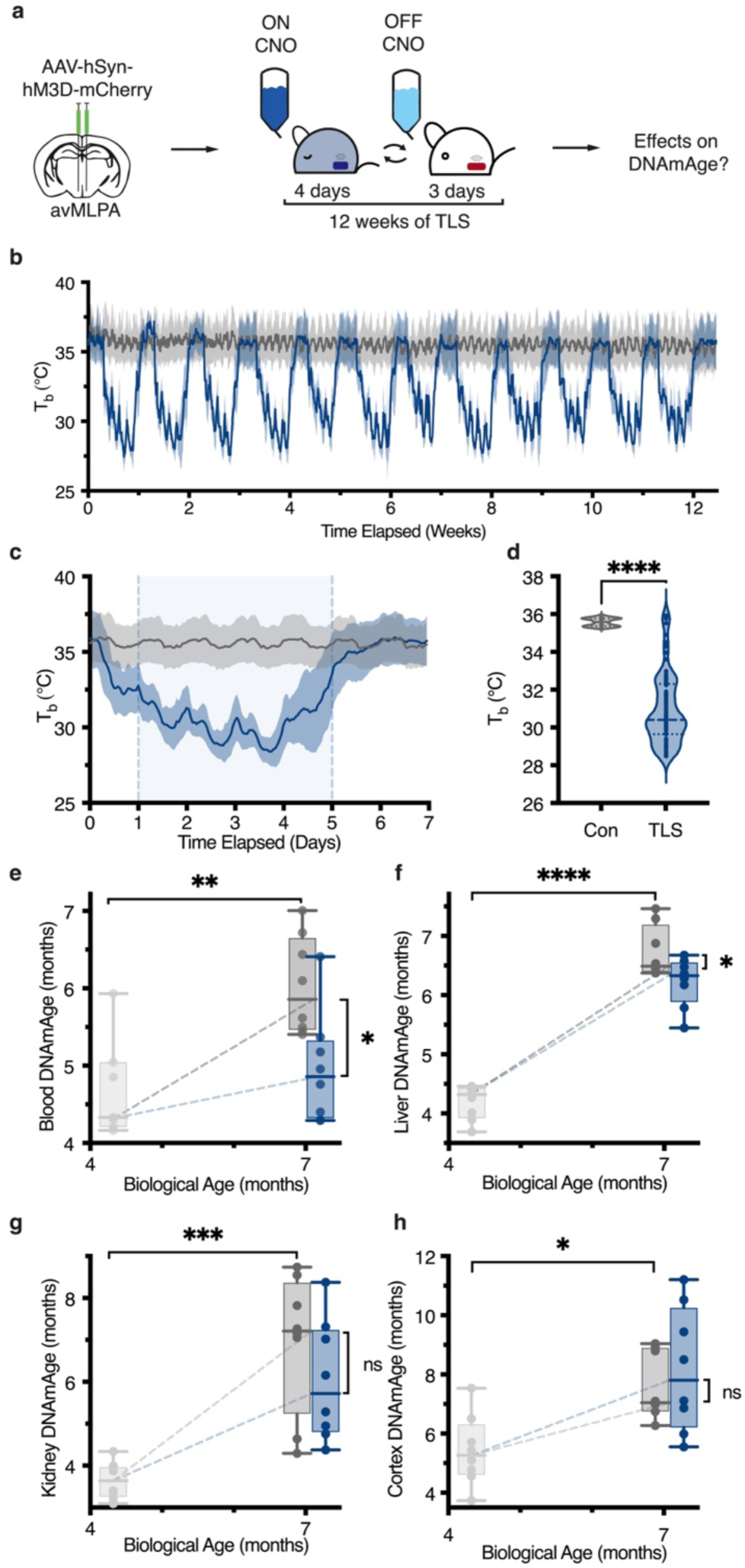
TLS slows epigenetic aging across multiple tissues. **a**, Schematic of long-term TLS induction through repeated CNO administration **b**, T_b_ of TLS (blue) and control (grey) mice over 12 weeks (*n* = 8). Line represents mean; shading denotes ± SD. **c**, Aggregate plot of T_b_ of TLS and control mice over 12-week experiment displayed over a 1-week interval. CNO administration marked by light blue shading between days 1-5. Lines and shading as in **b**. **d**, TLS mice had significantly lower average T_b_ per week (30.96 ± 0.19 °C) than control mice (35.58 ± 0.02 °C) while on CNO as determined by T-Test (*\*\*\*\* P* < 0.0001). Data reported as mean ± SEM. Data plotted as average T_b_ of individuals per week while on CNO. **e-h**, Epigenetic age (DNAmAge) as measured using tissue-specific epigenetic clocks of age-matched mice before the experiment (T0, *n* = 8) and after 3 months (control, *n* = 8, TLS *n* = 8). Data plotted as box plots from min-to-max with line at median. We found significant differences in epigenetic age between T0 and control mice across all tissues as measured by one-way ANOVA adjusted for multiple comparisons by Tukey’s HSD, validating the ability of epigenetic clocks to capture age-related changes over the time period (blood, \*\**P* = 0.0025, liver, \*\*\*\**P* < 0.0001, kidney, \*\*\**P* = 0.0001, cortex, \**P* = 0.0274). After 3 months, TLS mice had significantly lower epigenetic age in the blood (4.96 ± 0.24) (**e**) and liver (6.23 ± 0.14) (**f**) than control mice (blood = 6.03 ± 0.21, \**P* = 0.011, liver = 6.73 ± 0.14, \**P* = 0.043). We found no significant differences in the epigenetic age of the kidney (ns, *P* = 0.348) (**g**) and cortex (ns, *P* = 0.7438) (**h**) between control (kidney = 6.03 ± 0.47, cortex = 7.58 ± 0.38) and TLS mice (kidney = 6.94 ± 0.55, cortex = 8.15 ± 0.69) as measured by one-way ANOVA adjusted for multiple comparisons by Tukey’s HSD. Data reported as mean ± SEM.

### Long-term induction of TLS causes a cumulative and sustained decrease in blood epigenetic age and improves health span

As aging is a gradual and complex process, we induced TLS over a prolonged duration to better capture its effects on both epigenetic aging and age-related decline in physiological function. We serially measured blood epigenetic age in mice after 0, 3, 6, and 9 months of TLS (Fig. 3a). TLS had a linear, cumulative effect on blood epigenetic age: while at T0 TLS and control mice had equivalent blood epigenetic age, TLS mice appeared ∼3.0±0.6 months younger than control mice after 9 months of TLS (Fig. 3a-c). We used a linear regression to estimate the rate of epigenetic aging over 9 months of TLS, and found that TLS reduced the rate of blood epigenetic aging by 36.9% (F = 58.14, DFn = 1, DFd = 70, \*\*\*\**P* < 0.0001) (Fig. 3a). Longitudinal quantification of the rate of blood epigenetic aging across serial measurements from individual mice confirmed this finding; on average, individual TLS mice aged 38.9% slower than control mice (Fig. 3b). To test whether the pro-longevity effects of TLS are sustained, we measured blood epigenetic age in TLS mice 3 months after cessation of TLS (Fig. 3d). At this time, TLS mice still appeared ∼1.5 ±0.4 months younger than control mice (\*\*\**P* = 0.00013), indicating that TLS induces sustained epigenetic remodeling that was not rapidly reversed following exit from this state (Fig. 3e).

**Fig. 3.**
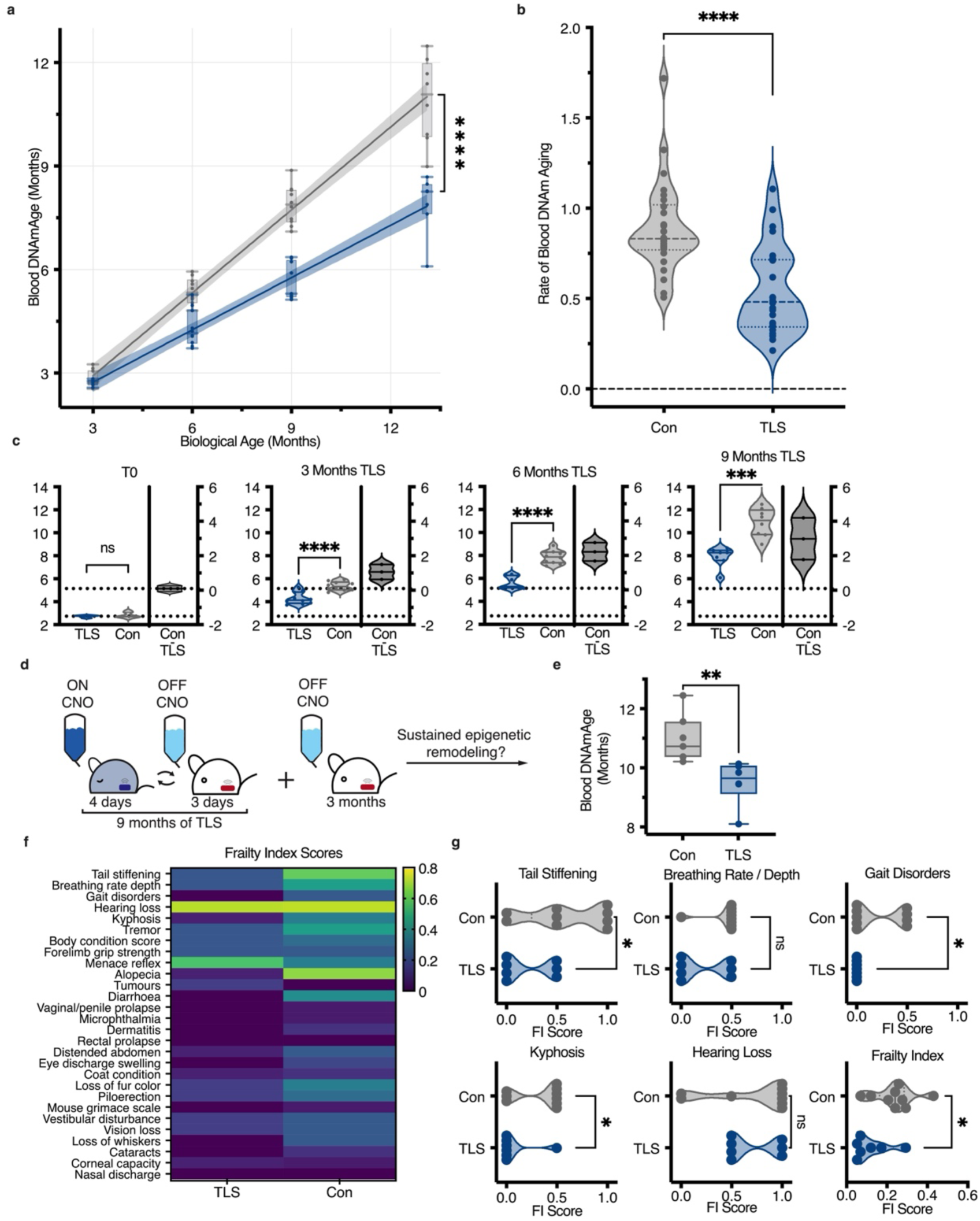
Long-term induction of TLS causes a cumulative and sustained decrease in blood epigenetic age and improves health span. **a**, Blood epigenetic age was serially measured every 3 months in control (grey) and TLS (blue) mice over 9 months. Data plotted as box plots from min-to-max with line at median. Simple linear regression was used to calculate the rate of blood epigenetic aging over 9 months; line represents regression, shading denotes 95% confidence interval (CI). TLS mice had a significantly slower rate of blood epigenetic aging 0.51 [0.46, 0.56] (r^2^ = 0.924, \*\*\*\**P* < 0.0001) than control mice 0.81 [0.74, 0.86] (r^2^ = 0.953, \*\*\*\**P* < 0.0001) (F = 58.14, DFn = 1, DFd = 70, \*\*\*\**P* < 0.0001) over 9 months of TLS. Data reported as mean with 95% CI in brackets. **b,** Quantification of the average rate of epigenetic aging measured every 3-months for 9-months across individual mice. TLS mice had a significantly slower rate of epigenetic aging (*n* = 26, 0.545 ± 0.047) than control mice (*n* = 26, 0.892 ± 0.05) (\*\*\*\**P* < 0.0001). Data reported as mean ± SEM. **c**, Estimation plots of the difference between means of TLS and control mice across measured timepoints. Data reported as mean ± SEM. Before treatment began (T0), control (2.83 ± 0.075) and TLS (2.74 ± 0.03) mice had equivalent blood epigenetic age (*n* = 10, ns, *P* = 0.252). After 3, 6, and 9 months of TLS, TLS mice had increasingly lower mean epigenetic age than controls (at 3 months, *n* = 10, TLS = 4.31 ± 0.168, Con = 5.37 ± 0.121, \*\*\*\**P* < 0.0001; at 6 months; TLS = 5.67 ± 0.180, Con = 7.87 ± 0.178, *n* = 9, \*\*\*\**P* < 0.0001, at 9 months, TLS = 7.90 ± 0.330, Con = 10.89 ± 0.434, *n* = 7, \*\*\**P* = 0.0001). **d,** Schematic of testing for sustained epigenetic remodeling. After 9 months of TLS, mice were off CNO for 3 months, after which time blood epigenetic age was measured again. **e,** Quantification of blood epigenetic age 3 months post-TLS. Data shown as box plots from min-to-max with line at median. TLS mice (9.51 ± 0.305) still had significantly younger blood epigenetic age than control mice (10.96 ± 0.305), (*n* = 6, \*\**P* = 0.0065). Data reported as mean ± SEM. **f**, Heatmap of the average scores on Frailty Index measurements of TLS and control mice after 9 months of TLS (*n* = 7). Frailty Index measurements are arranged in decreasing order of strongest correlation with age. **g**, Violin plots of FI scores of TLS mice and control mice on the five individual FI measurements that most strongly correlate with age and overall Frailty Index score. TLS mice scored significantly lower than control mice (*n* = 7) on tail stiffening (TLS = 0.214 ± 0.101, Con = 0.611 ± 0.139, \**P* = 0.0463), gait disorders (TLS = 0 ± 0, Con = 0.222 ± 0.088, \**P* = 0.043), and kyphosis (TLS = 0.071± 0.071, Con = 0.333 ± 0.083, \**P* = 0.037), as well as overall FI score, (TLS = 0.118 ± 0.034, Con = 0.238 ± 0.035, \**P* = 0.0290). Data reported as mean ± SEM.

Aging causes the accumulation of physical and physiological deficits over time. Composite measures of deficit accumulation, such as the mouse clinical frailty index (FI), serve as powerful tools to measure biological aging. Indeed, the mouse FI, is comprised of 31 non-invasive measurements, and is strongly correlated with and predictive of age and longevity^27,28^. To examine if TLS had an effect on functional measures of aging, we profiled mice using the FI assessment. After 9 months of TLS, TLS mice scored significantly lower on the FI assessment than age-matched controls (\**P* = 0.0290), indicating that they had an improved health span, and suggesting that they were functionally younger (Fig. 3f, Extended Data Fig. 4a, b). Notably, several individual FI parameters that most strongly correlate with age (r^2^ > 0.35*, P* < 1e^−30^)^27^ showed significant differences between TLS and control mice, including tail stiffening (\**P* = 0.0463), gait disorders *(*P =* 0.043), and kyphosis *(*P =* 0.037)^27^, further indicating that TLS slows age-related decline in physiological function (Fig. 3g). Taken together, these findings demonstrate that prolonged TLS slows molecular measures of aging and extends health span in mice.

### Decreased T_b_ mediates the rate of blood epigenetic aging during TLS

Hibernation and TLS are complex states characterized by a host of physiological changes that have been heavily implicated in aging and longevity, most notably decreased metabolic rate (MR), long-term caloric restriction (CR), and decreased T_b_^29,30^. In natural states these factors are inextricably intertwined, rendering them nearly inseparable. Here, we harnessed our inducible model of TLS to isolate the effects of decreased metabolic rate, long-term caloric restriction, and decreased T_b_ on blood epigenetic aging.

Metabolic rate has long been thought to modulate aging, as first explained in the ‘rate-of-living’ theory, founded on the observation that smaller animals with faster metabolisms have shorter lifespans than larger animals with slower metabolisms^31–33^. Importantly, metabolic rate and T_b_ often change in parallel, making it difficult to establish a unique role for either mechanism underlying pro-longevity interventions, including natural hibernation. We sought to decouple changes in metabolism from changes in T_b_ during TLS by stimulating avMLPA neurons in mice housed at a thermoneutral temperature (T_a_= 32°C) so as to eliminate the stimulation-induced drop in T_b_ (Fig. 4a). We found that stimulation of the avMLPA in mice housed at thermoneutrality largely recapitulated the metabolic changes seen in TLS independent of large decreases in T_b_ (Fig. 4b). Most notably, mice at thermoneutrality exhibited a similar decrease in metabolic rate to that of mice during TLS (ns, *P* = 0.1819) (Fig. 4b).

**Fig. 4.**
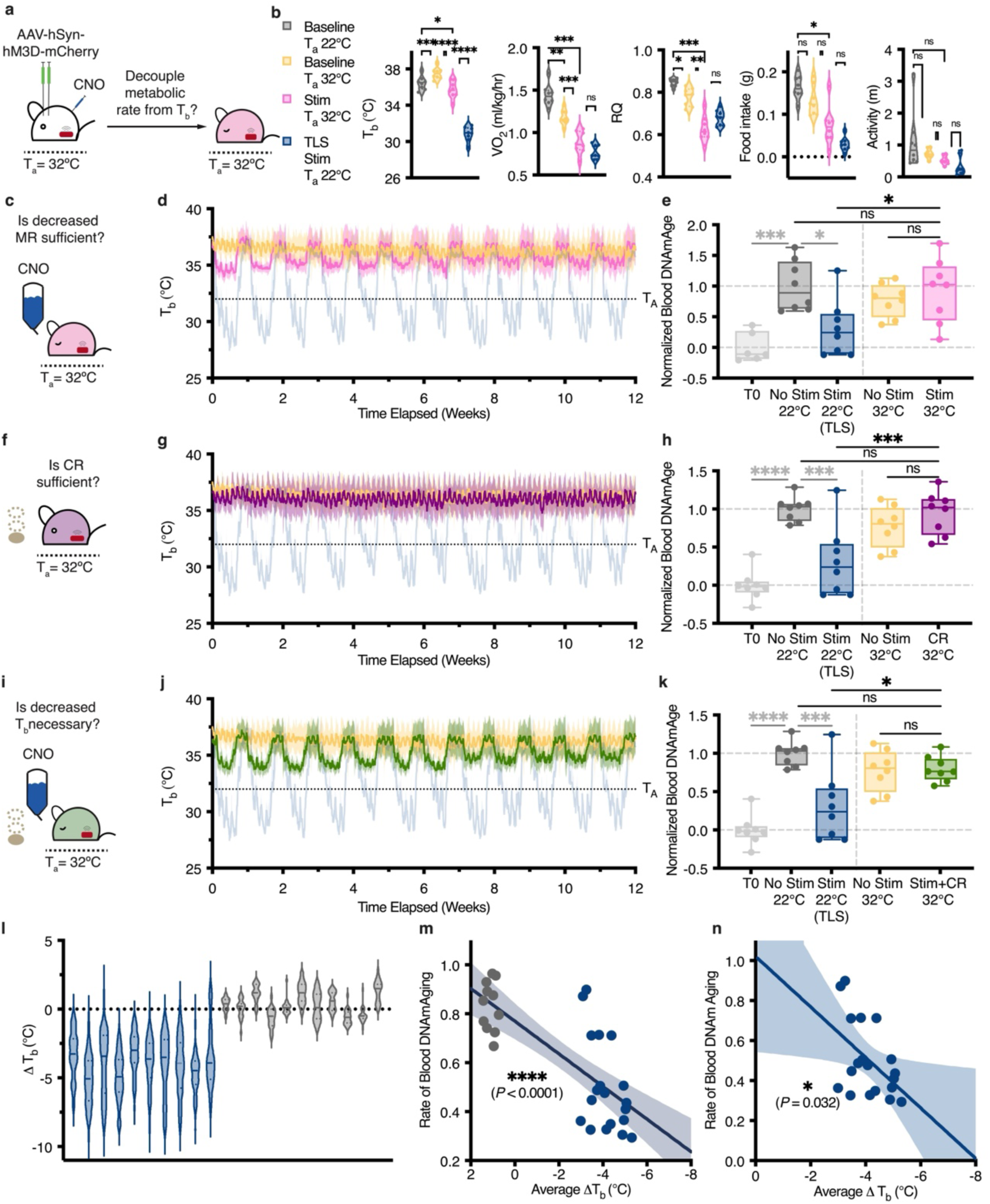
Decreased Tb mediates the rate of blood epigenetic aging during TLS. **a,** Schematic of stimulating avMLPA neurons while housing mice at thermoneutrality (Ta = 32°C) to decouple changes in metabolic rate from changes in temperature. **b,** Quantification of Tb, metabolic rate (VO2), RQ, food intake, and activity of mice at baseline and during stimulation of the avMLPA while housed at thermoneutrality (Ta = 32°C) as compared to mice at baseline and in TLS (Ta = 22°C). Data plotted as the average of each individual mouse. (T_b_ Baseline T_a_ 22°C = 36.44 ± 0.26, Baseline T_a_ 32°C = 37.50 ± 0.22, Stim T_a_ 32°C = 35.73 ± 0.32, TLS (Stim T_a_ 22°C) = 30.85 ± 0.28, \*\*\*\**P* < 0.0001) (VO_2_ Baseline T_a_ 22°C = 1.44 ± 0.04, Baseline T_a_ 32°C = 1.17 ± 0.03, Stim T_a_ 32°C = 0.88 ± 0.05, TLS (Stim T_a_ 22°C) = 0.78 ± 0.03, \*\*\*\**P* < 0.0001) (RQ Baseline T_a_ 22°C = 0.85 ± 0.01, Baseline T_a_ 32°C = 0.78 ± 0.02, Stim T_a_ 32°C = 0.63 ± 0.02, TLS (Stim T_a_ 22°C) = 0.68 ± 0.01, \*\*\*\**P* < 0.0001) (Activity Baseline T_a_ 22°C = 1.11 ± 0.33, Baseline T_a_ 32°C = 0.73 ± 0.06, Stim T_a_ 32°C = 0.52 ± 0.05, TLS (Stim T_a_ 22°C) = 0.33 ± 0.10, ns, *P* = 0.0706) (Food Intake Baseline T_a_ 22°C = 0.16 ± 0.01, Baseline T_a_ 32°C = 0.13 ± 0.01, Stim T_a_ 32°C = 0.073 ± 0.02, TLS (Stim T_a_ 22°C) = 0.03 ± 0.01, \*\*\*\**P* < 0.0001). **c,** Schematic of experimental design to determine sufficiency of decreased metabolic rate for effects on epigenetic aging. **d,** Tb over 12-week experiment of mice undergoing stimulation of avMLPA neurons while housed at thermoneutrality *(Stim 32*°*C*, *n* = 7) plotted in pink and control mice housed at thermoneutrality (*Con 32*°*C*, *n* = 8) plotted in orange. Tb of *TLS* mice shown in blue for reference (as previously shown in Fig. 2b); lines represent means, shading denotes SD. **e,** To compare changes in epigenetic aging across experiments, blood DNAmAge was normalized to *T0* = 0 ± 0.098 and *No Stim 22°C* = 1 ± 0.144 (shown for reference, previously shown in Fig. 2b); data plotted as box-plots min-to-max with line at median, data reported as mean ± SEM. Significance determined by one-way ANOVA adjusted for multiple comparisons by Tukey’s HSD. *Stim 32*°*C* (0.927 ± 0.184) had similar blood epigenetic age to *No Stim 32*°*C* mice (0.771 ± 0.095) (ns, *P* = 0.7848) and *No Stim 22°C* mice (1.00 ± 0.144) (ns, *P* = 0.9961). *Stim 32°C* had higher blood epigenetic age than TLS mice (previously shown in Fig. 2b); (0.306 ± 0.163) (\**P* = 0.0309). **f,** Schematic of experimental design to determine sufficiency of caloric restriction for effects on epigenetic aging. **g,** Tb over 12-week experiment of mice pair-fed with TLS mice while housed at thermoneutrality (*CR 32*°*C*, *n* = 8) plotted in purple. Tb of *No Stim* 32°C mice (orange) and TLS mice (blue) shown for reference (as previously shown in (**d**)). Lines and shading as in (**d**). **h,** Quantification of normalized blood DNAmAge of *CR 32°C* mice. Normalization performed as in (**e**) (*T0* = 0 ± 0.069, *No Stim 22°C* = 1± 0.057). *CR 32°C* mice had similar blood epigenetic age to *No Stim 22°C* mice (ns, *P* = 0.9990) and *No Stim 32°C* mice (ns, *P* = 0.7893), and significantly higher epigenetic age than *TLS* mice (\*\*\**P* = 0.0004) (shown for reference, as previously shown in Fig. 2b). **i,** Schematic of experimental design to determine the necessity of decreased Tb for effects on epigenetic aging. **j,** Tb over 12-week experiment of mice pair-fed with TLS mice and undergoing avMLPA stimulation housed at 32°C *(Stim +CR 32*°*C*, *n* = 8) plotted in green. Tb of *No Stim 32°C* mice (orange) and TLS mice (blue) shown for reference (as shown in (**d**)). Lines and shading as in (**h**). **k,** Quantification of normalized blood DNAmAge of *Stim +CR 32*°*C* mice. Normalization performed as in (**e**). *Stim +CR 32*°*C* mice (0.798 ± 0.058) had similar blood DNAmAge to *No Stim 32*°*C* mice (ns, *P* > 0.9999) and *No Stim 22°C* mice (ns, *P* = 0.6854). *Stim+CR 32°C* mice had significantly higher blood epigenetic age than *TLS* mice (\**P* = 0.0108) (shown for reference, as previously shown in Fig. 2b). **l,** ι1Tb of individual control and TLS mice while on CNO over 9 months (mice previously shown in Fig. 3a). **m,** Correlation between the average ι1Tb and the rate of blood DNAmAging in individual *Con 22°C* (grey) and *TLS* (blue) mice over 9 months (r^2^ = 0.587, \*\*\*\**P* < 0.0001); line represents simple linear regression, shading denotes 95% CI **n,** Correlation between the average ι1Tb and the rate of blood DNAmAging in *TLS* mice over 9 months (r^2^ = 0.243, \**P* = 0.032). Data shown as in (**m**).

To test whether a decrease in metabolic rate is sufficient to slow blood epigenetic aging, we repeatedly stimulated avMLPA neurons over 3-months in mice housed at thermoneutrality (*Stim 32°C*), mimicking the pattern of long-term TLS (Fig. 4c). *Stim 32°C* mice had equivalent average T_b_ to control mice housed at 22°C (*Con 22°C*), indicating that we were able to decouple changes in metabolic rate from changes in T_b_ (ns, *P* = 0.9941) (Fig. 4d, Extended Data Fig. 5a, e-f). After 3 months, we measured blood epigenetic age and found that *Stim 32°C* mice had equivalent blood epigenetic age to both control mice housed at 22°C (*No Stim 22°C*) (ns, *P* = 0.9961) and control mice housed at 32°C (*No Stim 32°C*) (ns, *P* = 0.7848) (Fig. 4e). Importantly, *Stim 32°C* mice had significantly higher blood epigenetic age than TLS mice (\**P* = 0.0309) (Fig. 4e), indicating that decreased metabolic rate alone is not sufficient to reproduce the pro-longevity effects of TLS on blood epigenetic aging.

Having found that decreased metabolic rate was insufficient to recapitulate the effects of TLS on blood epigenetic aging, we tested whether caloric restriction underlies the pro-longevity effects of TLS. Caloric restriction is one of the most well-known and effective anti-aging interventions shown to extend lifespan across species^34^. Importantly, caloric restriction in mice results in time-restricted feeding and long fasting intervals during which mice routinely enter torpor, as characterized by decreases in T_b_^35^. To isolate the effects of caloric restriction independent of drops in T_b_ during fasting-induced daily torpor, mice were housed at thermoneutrality and pair-fed to match food intake in TLS mice (0.95 g per day) (*CR 32°C*) (Fig. 4f, Extended Data Fig. 5a-d, g-h). *CR 32°C* mice had similar average T_b_ to *No Stim 22°C mice*, indicating that we were able to isolate the effects of caloric restriction from T_b_ (ns, *P* = 0.9030) (Fig. 4g, Extended Data Fig. 5a-b, g-h). After 3 months, *CR 32°C* mice had equivalent blood epigenetic age to both *No Stim 22°C* mice (ns, *P* = 0.9990) and *No Stim 32°C* mice (ns, *P* = 0.7893) (Fig. 4h). *CR 32°C* mice had significantly higher blood epigenetic age than TLS mice (\*\*\**P* = 0.0004) (Fig. 4h), suggesting that caloric restriction in the absence of T_b_ decreases during natural fasting-induced daily torpor is not sufficient to recapitulate the effects of TLS on blood epigenetic aging.

Given that caloric restriction and decreased metabolic rate on their own were insufficient to slow blood epigenetic aging, we hypothesized that decreased T_b_ was necessary for the pro-longevity effects of TLS. To isolate the effect of decreased T_b_ during TLS on blood epigenetic aging, we housed mice at thermoneutrality and both stimulated avMLPA neurons and pair-fed them with TLS mice (*Stim + CR 32°C*) (Fig. 4i, Extended Data Fig. 5a-d, i-j). *Stim +CR 32°C* mice thus exhibited both a decreased metabolic rate and long-term caloric restriction independent of large decreases in T_b_. *Stim+CR 32°C* mice had equivalent average T_b_ to *No Stim 22°C* mice (ns, *P* = 0.9915), indicating that we were able to blunt large decreases in T_b_ characteristic of TLS (Fig. 4j, Extended Data Fig. 5a, i). After 3 months, we measured blood epigenetic age and found that *Stim + CR 32°C* mice had equivalent blood epigenetic age to both *No Stim 22°C* mice (ns, *P* = 0.6854) and *No Stim 32°C* mice (ns, *P* > 0.9999) (Fig. 4k). Importantly, *Stim+CR 32°C* mice had significantly higher blood epigenetic age than TLS mice, (\**P* = 0.0108), indicating that despite having both a decreased metabolic rate and undergoing long-term caloric restriction, *Stim+CR 32°C* mice do not recapitulate the effects of TLS on blood epigenetic aging (Fig. 4k). Differential methylation analysis further supported the temperature-dependence of epigenetic remodeling during TLS (Extended Data Fig. 6a-l). Taken together, our results demonstrate the insufficiency of caloric restriction and decreased metabolic rate, both alone and when combined, to recapitulate the effects of TLS on blood epigenetic age, thus pointing to decreased T_b_ as necessary for the pro-longevity effects of TLS.

In light of the necessity of decreased T_b_ to capture the effects of TLS on blood epigenetic aging, we took advantage of the variability in the depth of TLS across individual animals by comparing the rate of blood epigenetic aging to the average decrease in T_b_ while on CNO over 9 months of TLS (Fig. 4l-n). This analysis revealed a significant correlation (r^2^ = 0.5869, **** *P* < 0.0001) between the average decrease in T_b_ and the rate of epigenetic aging across control and TLS mice (Fig. 4m). This correlation held significance in TLS mice alone (r^2^ = 0.2428, \**P* = 0.0321), further supporting the hypothesis that decreases in T_b_ mediate the effect of TLS on blood epigenetic aging (Fig. 4n). In summary, we show that long-term induction of a torpor-like state (TLS) through targeted stimulation of the avMLPA has pro-longevity effects on both functional and epigenetic measures of aging. While not possible in natural torpor or hibernation, we took advantage of the controllable nature of TLS to show that neither caloric restriction nor lower metabolic rate are sufficient and that instead decreased core body temperature is necessary for torpor-like states to slow blood epigenetic aging.

## Discussion

Our study examines the pro-longevity effects of hypothermic and hypometabolic states, such as torpor and hibernation, which are extreme physiological adaptations of homeotherms. While there are well-established links between hibernation, composed of multiple bouts of torpor, and longevity, how torpor and hibernation affect aging remains largely unexplored due to difficulties in studying natural hibernators. We demonstrate that the activity of a spatially defined neuronal population in the avMLPA, which has previously been identified as a torpor-regulating brain region, is sufficient to induce a torpor like state (TLS) in mice. TLS shares key metabolic and thermoregulatory features with both fasting-induced daily torpor and hibernation. We harnessed our model of TLS to explore the effects of torpor on aging and found that, similar to natural hibernation, TLS slows blood epigenetic aging. The effects of TLS on blood epigenetic age are both cumulative and sustained, agreeing with work performed in natural hibernators that found that the length of time spent in hibernation positively correlates with longevity, as measured by lifespan^36^. Importantly, long-term induction of TLS increased health span, measured using the mouse FI assessment, indicating that TLS has systemic and functional effects on aging processes.

We went on to leverage the controllable nature of TLS to isolate the effects of decreased metabolic rate, long-term caloric restriction, and decreased T_b_ on blood epigenetic aging. We found that a decrease in T_b_ is necessary to slow blood epigenetic aging during TLS, and that the rate of blood epigenetic aging is mediated by the degree of decrease in T_b_. Neither a decrease in metabolic rate nor caloric restriction, alone or combined, were sufficient to recapitulate the prolongevity effects of TLS on blood epigenetic age. Our finding that decreased T_b_ mediates the prolongevity effects of TLS agrees with recent work showing that T_b_ is a more important modulator of lifespan than metabolic rate^30^. Similarly, our finding that caloric restriction independent of a decrease in T_b_ does not slow blood epigenetic aging agrees with the previous finding that housing mice at thermoneutrality blunts the pro-longevity effects of CR^37^. Recent work on caloric restriction has shown that the lifespan extension effects of CR are primarily mediated by fasting intervals of >12 h during which mice likely enter torpor, reaffirming the possibility that torpor and decreased core body temperature play a central role in CR-mediated effects on longevity ^38^. Notably, decreases in T_b_ have previously been correlated with increased lifespan in transgenic mice ^29^. Our finding that decreased T_b_ is necessary to slow blood epigenetic aging during TLS further supports the growing body of evidence that T_b_ is an important mediator of aging processes.

It is important to recognize that torpor and hibernation are complex behaviors involving profound physiological changes that are not fully recapitulated in our model. In light of the necessity of decreased T_b_ for the effects of TLS on blood epigenetic aging, and the correlation between the depth of TLS and the rate of blood epigenetic aging, perhaps the most salient difference between TLS and natural hibernation is that mice in TLS exhibit shallower drops in T_b_ than some natural hibernators, who drop their T_b_ as low as 4°C. While TLS captures meaningful changes in both epigenetic aging and health span, TLS may not reproduce pro-longevity effects that might occur from the more extreme drops of T_b_ in natural hibernators. Future studies in natural hibernators should address the role of extreme drops in T_b_ in torpor-mediated increases in longevity.

Uncovering tissue-specific aging processes is critical to understanding the age-related decline of different organs and diverse cell types. We found that while TLS has systemic effects on functional measures of aging, TLS exerts tissue-specific effects on epigenetic aging whereby induction of TLS had greater effects on tissues with higher rates of basal cell turnover, most notably the blood. It is well known that temperature is a regulator of mammalian cell cycle, whereby moderate hypothermia arrests mitosis^39,40^. Our work showing the temperature-dependence of blood epigenetic aging coupled with the observation that there is a correlation between the rate of basal cell turnover in profiled tissues and the effect size of TLS on epigenetic aging offers a working framework for the observed tissue-specific effects of torpor-like states on aging that future studies should further address.

## Methods

### Mice

Experiments performed at Harvard Medical School were approved by the National Institute of Health and Harvard Medical School Institutional Animal Care and Use Committee. Experiments performed at the Whitehead Institute at the Massachusetts Institute of Technology were approved by the National Institute of Health and the Division of Comparative Medicine and the Committee on Animal Care. All experiments followed ethical guidelines described in the US National Institutes of Health Guide for the Care and Use of Laboratory Animals and were performed using adult C57BL/6J mice (The Jackson Laboratory, Stock 000664). Unless otherwise noted, all mice were group-housed at 22°C under a standard 12 h light/dark cycle and fed ad libitum. No statistical methods were used to predetermine sample size of groups and unless otherwise noted, all groups contained equal numbers of male and female mice randomly assigned to experimental groups before surgery.

### Telemetric monitoring of core body temperature

Mice were implanted with wireless probes (UID Cat# UCT-2112-, Temperature Microchip) in the intraperitoneal space. Mice were allowed to recover for at least four days before recording. To record core body temperature, cages were placed on the UID Mouse Matrix reader plate, allowing continuous and undisturbed tracking of core body temperature. Unless otherwise noted, core body temperature was logged every 30 s.

### Viral Constructs

pAAV-hSyn-mCherry (Addgene #114472-AAV8) and pAAV-hSyn-hM3D(Gq)-mCherry (Addgene #50474-AAV8) were obtained from Addgene. pAAV7-DJ-CMV-TeNT-P2A-GFP was prepared and obtained through the Stanford Viral Core. All viruses were diluted with PBS to a final concentration between 1-5 × 10^13^ genome copies per mL before stereotaxic delivery.

### Stereotactic viral injection

Mice were anesthetized via isoflurane inhalation and placed in a stereotactic headframe (Kopf Instruments). For all injections, coordinates AP +0.4MM, ML ± 0.5 MM, DV −0.5 mm relative to bregma were used to target the avMLPA. An air-based injection system was used to infuse the virus at a rate of approximately 100 nL min^-1^, and the needle was kept at the injection site for 5 minutes before withdrawing. For mice singly injected, 100-150 nL of virus was bilaterally injected. For mice co-injected, viruses were mixed before injection in a 1:1 ratio and 100-150 nL of virus was injected bilaterally.

### Fasting-induced daily torpor induction

Adult mice (*n* = 4F) were singly housed before food removal at an ambient temperature of 22°C. Food was removed at the beginning of the dark cycle, and initial bouts of torpor were seen in mice after approximately 10 hours of fasting. Natural torpor was defined by a T_b_ of < 35°C and lack of arousal. Food was returned to cages 24 h after the start of the fast.

### CNO administration

CNO solution was prepared by initially dissolving CNO hydrochloride (Sigma-Aldrich, SML2304) in H_2_O to a stock solution of 100 mM. For intraperitoneal injection of CNO, the stock solution was diluted with PBS to a final concentration of 0.6 mM, and approximately 250 μL was injected intraperitoneally per mouse for a final injection concentration of 2 mg kg^-1^. For continuous CNO administration, CNO was taken from a 100 mM stock solution and diluted in water to a final concentration between 0.01 mM – 0.04 mM.

### Image analysis

Mice were anesthetized via inhalation of isoflurane and euthanized by transcardial perfusion of 10 mL PBS followed by 10 ml of 4% paraformaldehyde (PFA). Brains were extracted, post-fixed overnight with 4% PFA at 4 °C and then embedded in PBS with 3% agarose. Brains were sliced on a vibratome (Leica VT1200S) into 50-μm coronal sections. Sections were imaged on an TissueFax SL Q 60um confocal disk slide scanner using a 10x 0.3 objective with extended focus and z-stacking.

### Metabolic Measurements

Mice (*n* = 4M, 4F) were implanted abdominally with telemetric temperature probes (Starr Life Science VV-EMITT-G2) and placed in the Sable Systems Promethion Core Metabolic System. Mice were given 24 h to acclimate before baseline recordings (2 days), followed by CNO water administration (5 days) and (3 days) recovery. Data was logged every 3 minutes. For analysis of natural torpor, mice were fasted as described previously. Six out of eight mice underwent natural fasting induced daily torpor bouts as defined by T_b_ <35°C and lack of arousal. For downstream analysis, the average value of metabolic parameters for each individual mouse was calculated across all torpor bouts.

### Statistical analysis of the thermoregulatory system

To analyze the thermoregulatory system during TLS and at baseline, parameters *G, T_set_* and *H* were estimated from observing T_b_ and VO_2_ across varying T_a_ (8, 16, 24°C) as previously described^10,15^. Mice (*n* = 11, 5F, 6M) were placed in Sable Systems Promethion Core Metabolic System for measurement of T_b_ and VO_2_. To record baseline and TLS data, mice were intraperitoneally injected with PBS and CNO, respectively, and T_a_ was lowered every 3 hours (24, 16, 8°C). The minimum VO_2_ and corresponding T_b_, theoretically corresponding to a metabolically stable state, of each individual mouse at each temperature were used in downstream analyses.

### Long-term TLS induction

Group-housed 16-week-old mice were either injected with pAAV-hSyn-mCherry (controls) or pAAV-hSyn-hM3D(Gq)-mCherry (TLS) and implanted with telemetric temperature probes. Mice were administered CNO water (0.01-0.04 mM) for 4 days, followed by 3 days of recovery over 12-weeks. Bodyweight was measured every two-weeks. Food-intake was measured twice a week – upon CNO administration and removal - allowing us to capture food consumption while on and off CNO.

### Tissue-processing

Mice were anesthetized via inhalation of isoflurane and euthanized by cervical dislocation. Blood was collected via cardiac puncture followed by transcardial perfusion of 10 mL PBS to wash tissues. Tissues were removed and washed in PBS twice before being flash frozen in liquid nitrogen. Following collection, tissues were stored at −80°C before further processing. For DNA extraction, DNeasy Blood & Tissue Kits were used (QIAGEN #69506). After extraction, DNA was stored at - 20°C until further processing. For RNA extraction, RNeasy Mini kits were used (QIAGEN #74104). After extraction, RNA was stored at −80°C until further processing.

### Methylation Analysis

DNA samples were normalized to between 12.5-25 ng/μL in 30 μL. DNA was bisulfite converted using the Zymo EZ DNA methylation kit (D5004). Bisulfite converted DNA was run on the Illumina Horvath Mammalian Methylation 320k Chip, which combines the Illumina Infinium Mouse Methylation 285k array with the mammalian methylation array, resulting in interrogation of > 285,000 CpGs specific to mice and 37,000 conserved mammalian loci^41^. The SeSaMe normalization method was used to estimate methylation levels (*β* values) for each CpG site.

### Epigenetic clock analysis

Our epigenetic clocks are based on the HorvathMammalMethylChip320. The data were generated and analyzed by the nonprofit Epigenetic Clock Development Foundation (Torrance, California) as previously described^21,22,24,41,42^. In our primary analysis (Fig. 2), we assessed the epigenetic ages of various mouse tissues by employing DNA methylation clocks tailored for blood, liver, kidney, and cerebral cortex. These mouse methylation clocks were developed based on distinct training datasets as reported by Mozhui (2022), with the corresponding R software code presented in the same study^42^. We also utilized the universal pan-mammalian epigenetic clocks (UniversalClock 2 and 3), as described in Nature Aging by A. Lu (2023) and depicted in Extended Figure 3^24^. These pan mammalian clocks leverage conserved cytosines to evaluate aging across mammalian species, highlighting epigenetic aging effects likely relevant to human aging. The R code of all of these epigenetic clocks can also be found in the MammalMethylClock R package^43^.

### Differential methylation analysis

Differential methylation analysis was performed using the SeSaMe pipeline. First, the DML function was used to identify differentially methylated probes between control and TLS mice. The DMR function was then used to align probes to the mouse genome (mm10) and find differentially methylated regions. The following thresholds for significance were used, an adjusted P-value < 10^-3^ and |ΔBeta| > 0.05.

### GREAT analysis

Gene enrichment and top biological processes analysis were performed using the Genomic Regions Enrichment of Annotations Tool (GREAT) v.4.0.4. In brief, differentially methylated regions were identified as described above. Only genomic regions with an adjusted p-value < 0.001 and |Δ*β*| > 0.05 were used for GREAT analysis with a basal plus extension of 5 kb proximal upstream, 1 kb proximal downstream, plus distal of up to 1 kb.

### Bulk RNA-seq library preparation and sequencing

Libraries were prepared using the NEBNext Ultra^TM^ II kit. In brief 500 ng of RNA per sample underwent Poly(A) selection before fragmentation and random priming followed by first and second strand cDNA synthesis. Following cDNA synthesis, NEBNext Multiplex Oligos for Illumina (E6440S) were ligated before PCR enrichment. All libraries were pooled to a final concentration of 1nM and sequenced on a NovaSeq 6000.

### Transcriptomic clock analysis

To assess the transcriptomic age (tAge) of liver, WAT, cortex and kidney derived from control mice and mice subjected to TLS, we applied a mouse multi-tissue gene expression clock of relative lifespan-adjusted age based on previously identified signatures of aging and lifespan-regulating interventions^44,45^. Genes with less than 5 reads in more than 87.5% of samples were filtered out, resulting in 17,695 expressed genes according to Entrez annotation. Filtered data was then passed to Relative Log Expression (RLE) normalization^46^. Normalized gene expression profiles were log-transformed and scaled. The missing values corresponding to clock genes not detected in the data were imputed with the precalculated average values. Samples from control 4-month-old animals were used as reference groups for every tissue. Differences between average tAges across the groups were assessed with one-sample ANOVA and adjusted for multiple comparisons using Tukey’s HSD.

### Differential gene expression

The following command was used to align reads to the mm10 reference mouse genome using STAR:

~~~
for fq1 in ‘/bin/ls *R1_001.fastq.gz’
do
 name=‘basename $fq1 | perl -pe ‘s/_R1_001.fastq.gz//’’
 fq2=‘basename $fq1 | perl -pe ‘s/R1_001.fastq.gz/R2_001.fastq.gz/’’
 echo Processing $name …
 # Slurm cluster call
 # echo sbatch --partition=20 --job-name=STAR --mem=32G --wrap “STAR –genomeDir /nfs/genomes/mouse_mm10_dec_11_no_random/STAR/GRCm38.102.canonical_over-hang_100 --sjdbScore 2 --runThreadN 8 --outSAMtype BAM SortedByCoordinate --read-FilesCommand zcat --readFilesIn $fq1 $fq2 --outFileNamePrefix ../STAR/$name.”
sbatch --partition=20 --job-name=STAR --mem=32G --wrap “STAR –genomeDir /nfs/genomes/mouse_mm10_dec_11_no_random/STAR/GRCm38.102.canonical_over-hang_100 --sjdbScore 2 --runThreadN 8 --outSAMtype BAM SortedByCoordinate --read-FilesCommand zcat --readFilesIn $fq1 $fq2 --outFileNamePrefix ../STAR/$name.”
done
~~~

After alignment, data was processed through the DESeq2 pipeline. In brief, data was separated by tissue-type and processed independently. DESeq() was run to normalize counts using contrast of treatment (TLS vs. Con). DESeq results were transformed to variance stabilized expression levels using the vst() function followed by log fold change shrinkage with the lfcShrink() function using apelgm. Data was visualized using the EnhancedVolcano() function with the following thresholds, (-log_10_(Pval) < 10^-6^ and Log_2_ fold change > 1.

### Clinical Frailty Index

Clinical frailty measurements were performed as previously described^27,28^. In brief, a blinded investigator scored mice on 31 noninvasive measures using a simple scale - mice were given a 0 if they showed no deficit, a 0.5 if they showed a mild deficit, and a 1 if they showed a severe deficit. All frailty measurements were performed when mice were off CNO and had recovered to euthermia. All bodyweight and temperature calculations were excluded from downstream frailty score analyses given the confounding effects of TLS on body temperature and bodyweight.

### Clamping T_a_

To blunt decreases in core body temperature, mice cages were placed at a thermoneutral temperature (32°C) in a Powers Scientific Incubator (IT54SD) for the duration of the experiment.

### Caloric Restriction

*CR 32°C* mice and *Stim + CR °32C* mice were pair-fed to TLS mice 4 days a week and fed ad libitum 3 days a week to isolate the effects of caloric restriction during TLS. In brief, *CR 32°C* mice, *Stim + CR °32C* mice, and *Con 32°C* mice were singly-housed at thermoneutral ambient temperature (32°C). We measured that while on CNO, TLS mice (1.17±0.1 g) ate on average 35% of what Con 22°C mice ate (3.31±0.14 g). Importantly, we found that while on CNO, *Stim 32°C* mice (2.30±0.12 g) ate on average 70% of what *Con 32°C* mice ate (3.25±0.22 g). When pair-feeding *CR 32°C* mice and *Stim + CR °32C* mice to TLS mice, we accounted for the difference in basal metabolic rates between mice housed at 32°C vs. 22°C, and thus fed mice 1±0.1g of food per day (roughly equivalent to 35% of the daily food intake of *Con 32°C* mice). While on CNO, *CR 32°C* mice and *Stim + CR °32C* were fed daily at the beginning of the dark cycle. While off CNO, *CR 32°C* mice and *Stim+CR°32C* mice were fed ad libitum. Food intake of all mice was measured twice weekly, once at the beginning of CNO treatment, and once at the end.

## Author Contributions

L.J. and S.Hrvatin conceived and designed the study. L.J. designed, performed, and analyzed experiments. A.L.P designed and performed experiments. A.T. performed transcriptomic clock analyses (tAge). M.A. designed and performed experiments. J.R., E.R., A.M., S.S., and H.Y. contributed to experiments. R.B. and S.Horvath applied epigenetic clocks to methylation data to generate DNAmAge estimates. V.N.G., V.J., and E.C.G. advised on the study. L.J., S.Hrvatin, and E.C.G wrote the manuscript. L.J., E.C.G., and S.Hrvatin. obtained funding for the research. All authors approved and reviewed the manuscript.

## Acknowledgements

We thank the UCLA Neuroscience Genomics Core (UNGC) for processing DNA samples for methylation analysis. We thank Michael E. Greenberg for providing feedback on the study. We thank the Clock Foundation for applying epigenetic clocks to generate DNAmAge estimates. We thank Beth Israel Deaconess Medical Center Metabolic Core (BIDMC) for performing metabolic experiments. We thank the Sinclair lab for teaching us how to perform frailty index measurements. This project was supported by a Longevity Impetus Grants from Norn Group to S.Hrvatin and V.N.G, the National Institute of Health grant 1DP2DK136123-01 to S.Hrvatin, and the National Institute of Aging grant AG067782 to V.N.G.

## Competing Interests

S. Horvath and R.T. Brooke are founders of the non-profit Epigenetic Clock Development Foundation, which licenses several patents from UC Regents including a patent on the mammalian methylation array platform. These patents list S.Horvath as inventor. V.G.S. serves as an advisor to and/or has equity in Branch Biosciences, Ensoma, and Cellarity, all unrelated to this work.

## Extended Data

**Extended Data Fig. 1.**
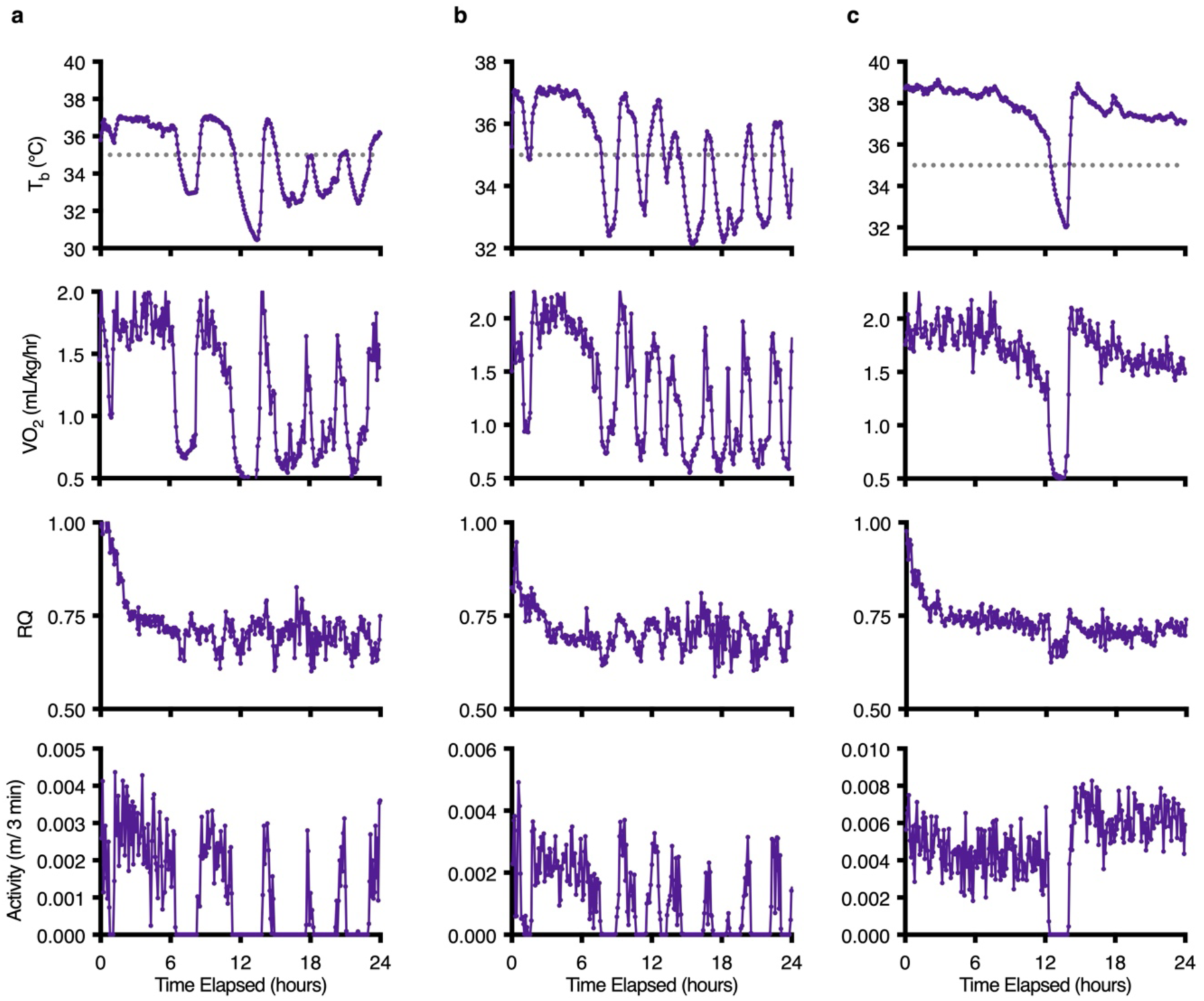
Metabolic characterization of natural fasting-induced daily torpor bouts. **a-f,** T_b_, VO_2_, RQ, and activity as measured by the Prometheon Metabolic System in 3 representative individual mice over a 24-hour fasting interval (food was removed at time 0). 6/8 mice underwent natural fasting-induced daily torpor bouts during the fasting interval as defined by T_b_ < 35°C and lack of arousal (threshold T_b_ for torpor entry marked by dotted grey line on T_b_ graph).

**Extended Data Fig. 2.**
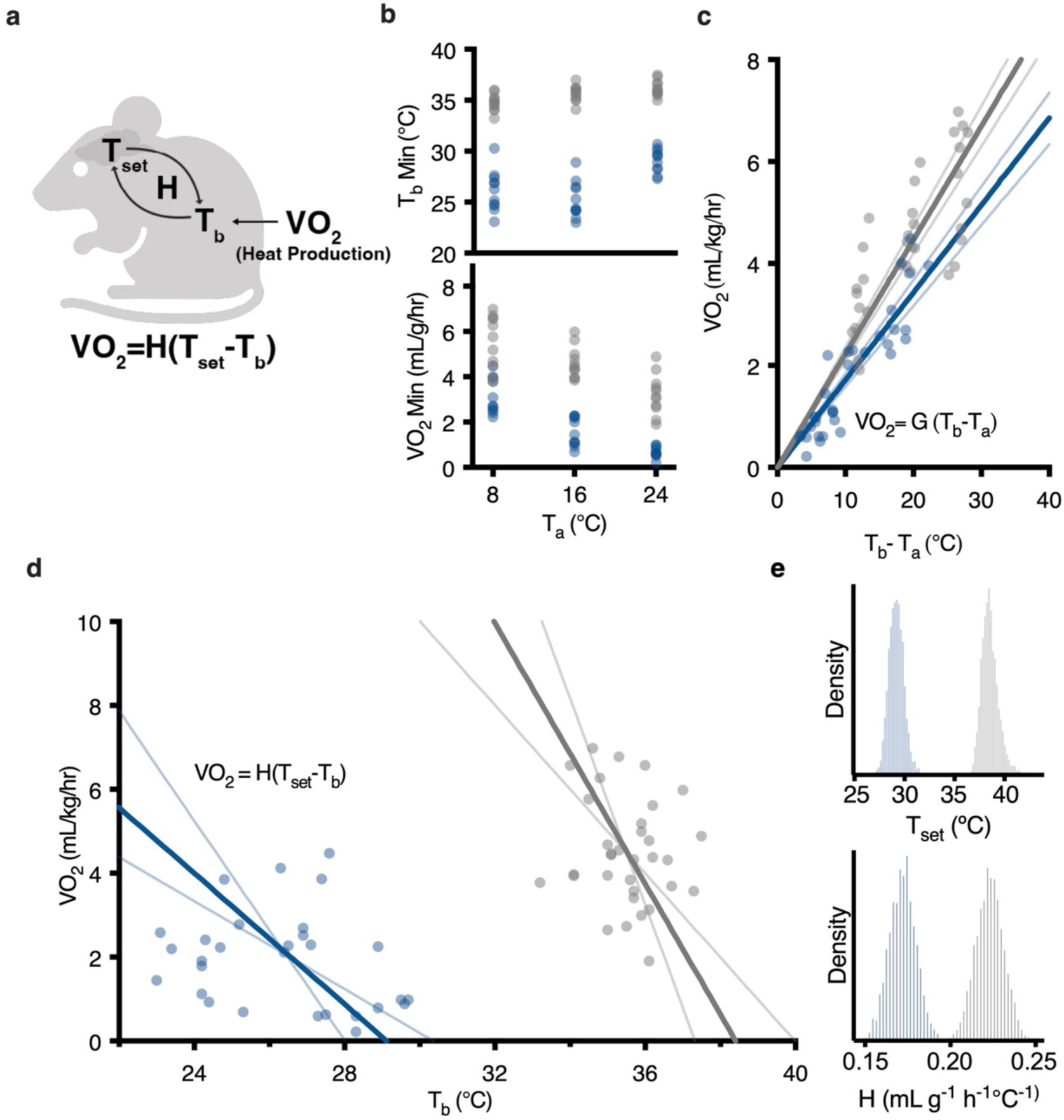
The thermoregulatory system during TLS. **a**, Schematic of the thermoregulatory model. **b**, Minimum T_b_ and VO_2_ across ambient temperatures (8, 16, 24 °C) at baseline and during TLS (*n* = 11). **c**, The relationship between VO_2_ and T_b_ - T_a_. The slope of the curve denotes the median value of *G*, the heat conductance, at baseline (0.233 ml g^-1^ h^-1^ °C^-1^) and during TLS (0.172 mL g^-1^ h^-1^ °C^-1^), thin lines denote the 89% highest posterior density interval (HPDI) of *G* at baseline [0.210, 0.236] mL g^-1^ h^-1^ °C^-1^ and during TLS [0.159, 0.184] mL g^-1^ h^-1^ °C^-1^, and dots represent data **d**, Relationship between T_b_ and VO_2_ across varying T_a_. The negative slope denotes the median value of *H* at baseline (1.555 g^-1^ h^-1^°C^-1^) and during TLS (0.780 g^-1^ h^-1^°C^-1^). The X-intercept, where VO_2_ = 0, represents the median theoretical set point temperature (T_set_) at baseline (38.414°C) and during TLS (29.13°C), thin lines represent the 89% HPDI of both *H* [1.00, 2.46] g^-1^ h^-1^°C^-1^ at baseline and [0.52, 1.31] g^-1^ h^-1^°C^-1^ during TLS and *T_set_* [37.323, 40.0]°C at baseline and 29.13 [28.02, 30.36]°C during TLS), dots represent data. **e**, Distribution of estimated T_set_ and H both at baseline and during TLS.

**Extended Data Fig. 3.**
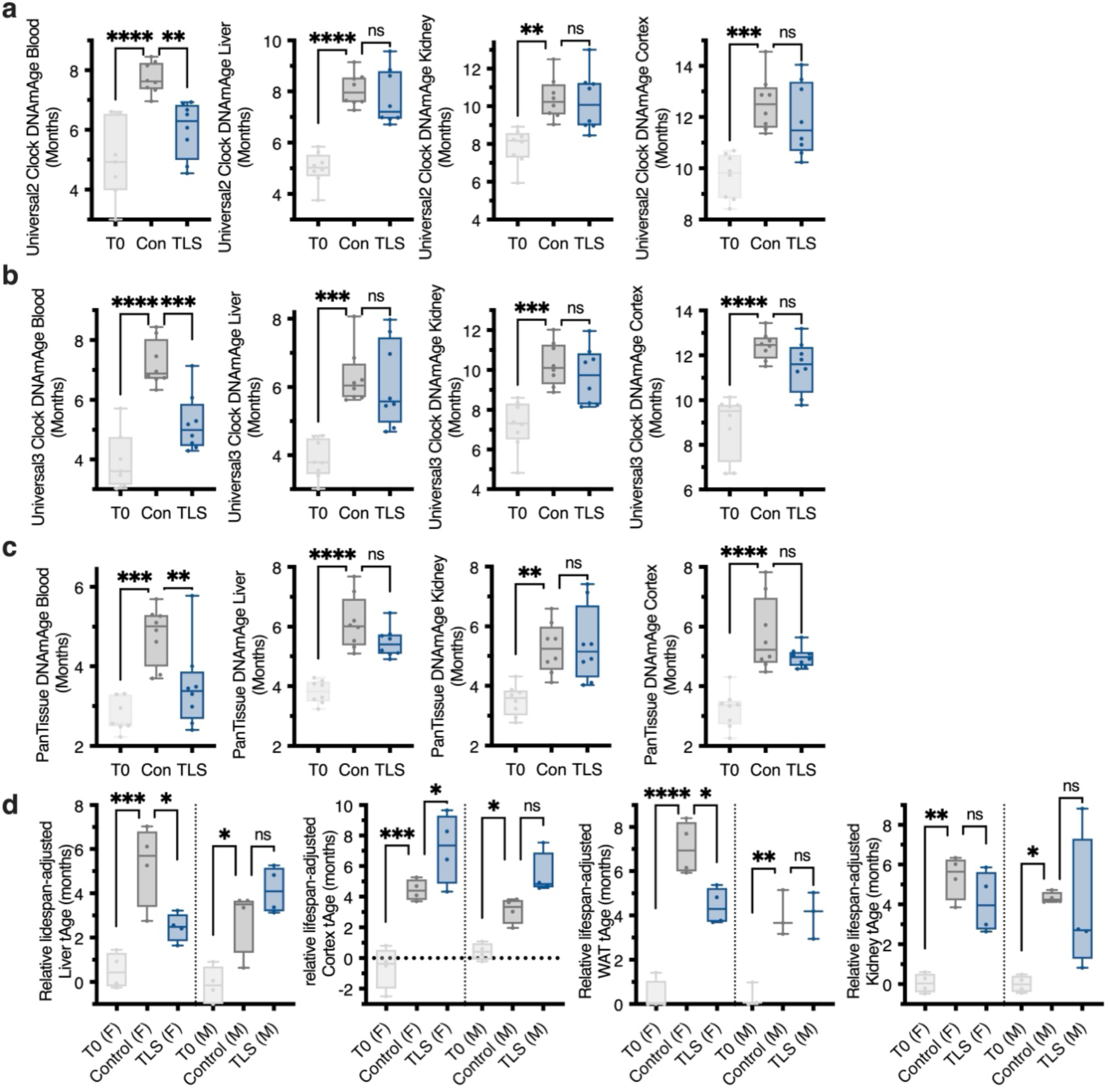
Additional epigenetic and transcriptomic clock analyses. **a,** DNAmAge across tissues as calculated using the Universal2 Epigenetic Clock. Data reported as mean ± SEM. Significance determined by one-way ANOVA adjusted for multiple comparisons by Tukey’s HSD. In the blood, TLS mice (6.01 ± 0.33) had significantly lower DNAmAge than control mice (7.34 ± 0.18) (\*\**P* = 0.0015). In the liver, kidney, and cortex, TLS mice (liver = 7.76 ± 0.39, kidney = 10.25 ± 0.56, cortex = 11.92 ± 0.51) had comparable DNAmAge to control mice (liver = 8.08 ± 0.23, kidney = 10.40 ± 0.39, cortex = 12.54 ± 0.38) (liver = ns, *P* = 0.7240, kidney = ns, *P* = 0.9705, cortex = ns, *P* = 0.5352). **b,** DNAmAge across tissues as calculated using the Universal3 Epigenetic Clock. Data reported as mean ± SEM. Significance determined by one-way ANOVA adjusted for multiple comparisons by Tukey’s HSD. In the blood, TLS mice (5.22 ± 0.34) had significantly lower DNAmAge than control mice (7.21 ± 0.27) (*** *P* = 0.0008). In the liver, kidney, and cortex, TLS mice (liver = 6.08 ± 0.45, kidney = 9.71 ± 0.51, cortex = 11.50 ± 0.41) had equivalent DNAmAge to control mice (liver = 6.28 ± 0.29, kidney = 10.31 ± 0.39, cortex = 12.41 ± 0.22) (liver = ns, *P* = 0.7240, kidney = ns, *P* = 0.6152, cortex = ns, *P* = 0.2434). **c,** DNAmAge across tissues as calculated using the PanTissue Epigenetic Clock. Data reported as mean ± SEM. Significance determined by one-way ANOVA adjusted for multiple comparisons by Tukey’s HSD. In the blood, TLS mice (3.5 ± 0.37) had significantly lower DNAmAge than control mice (4.80 ± 0.26) (\*\**P* = 0.0099). In the liver, kidney, and cortex, TLS mice (liver = 5.478 ± 0.18, kidney = 5.40 ± 0.45, cortex = 4.98 ± 0.12) had equivalent DNAmAge to control mice (liver = 6.13 ± 0.32, kidney = 5.27 ± 0.30, cortex = 5.72 ± 0.44) (liver = ns, *P* = 0.1484, kidney = ns, *P* = 0.959, cortex = ns, *P* = 0.1997). **d,** Transcriptomic clock analysis across tissues. Data reported as mean. Significance determined by one-way ANOVA adjusted for multiple comparisons by Tukey’s HSD. In the liver, there were significant differences between T0 and control mice in females (F) and males (M) (F T0 = 0.52, F control = 5.29, \*\*\**P* = 0.0003; M T0 = −0.24, M control = 2.835, \**P* = 0.022). There were significant differences between control and TLS mice in females (F control =5.29, F TLS = 2.46, *\*P =* 0.039), but not males (M control =2.835, M TLS = 4.15, ns, *P* = 0.649). In the cortex there were significant differences between T0 and control mice in both males and females (F T0 = −0.60, F control = 4.44, \*\*\**P* = 0.0002) (M T0= 0.43, M control = 3.12, \**P* = 0.0416). There were significant differences between control and TLS mice in females (F control = 4.44, F TLS = 7.17, \**P* = 0.0387), but not in males (M control = 3.12, M TLS = 5.45, ns, *P* = 0.0926). In the WAT, there were significant differences between T0 and control mice in both females and males (F T0 = 0.17, F control = 7.05, *\*\*\*\*P* < 0.0001) (M T0 = 0.22, M control = 4.00, \*\**P* = 0.0025). There were significant differences between control and TLS mice in females (F control = 7.05, F TLS = 4.42, \**P* = 0.0153), but not in males (M control = 4.00, M TLS = 4.06, ns, *P* > 0.9999). In the kidney, there were significant differences between T0 and control mice in both females (F T0 = 0.05, F control = 5.37, \*\**P* = 0.0027) and males (M T0 = 0.01, M control = 4.33, \**P* = 0.0166). There were no significant differences between control and TLS mice in females or males (F control = 5.37, F TLS = 4.11, ns, *P* = 0.8792; M control = 4.33, M TLS = 3.76, ns, *P* = 0.9960).

**Extended Data Fig. 4.**
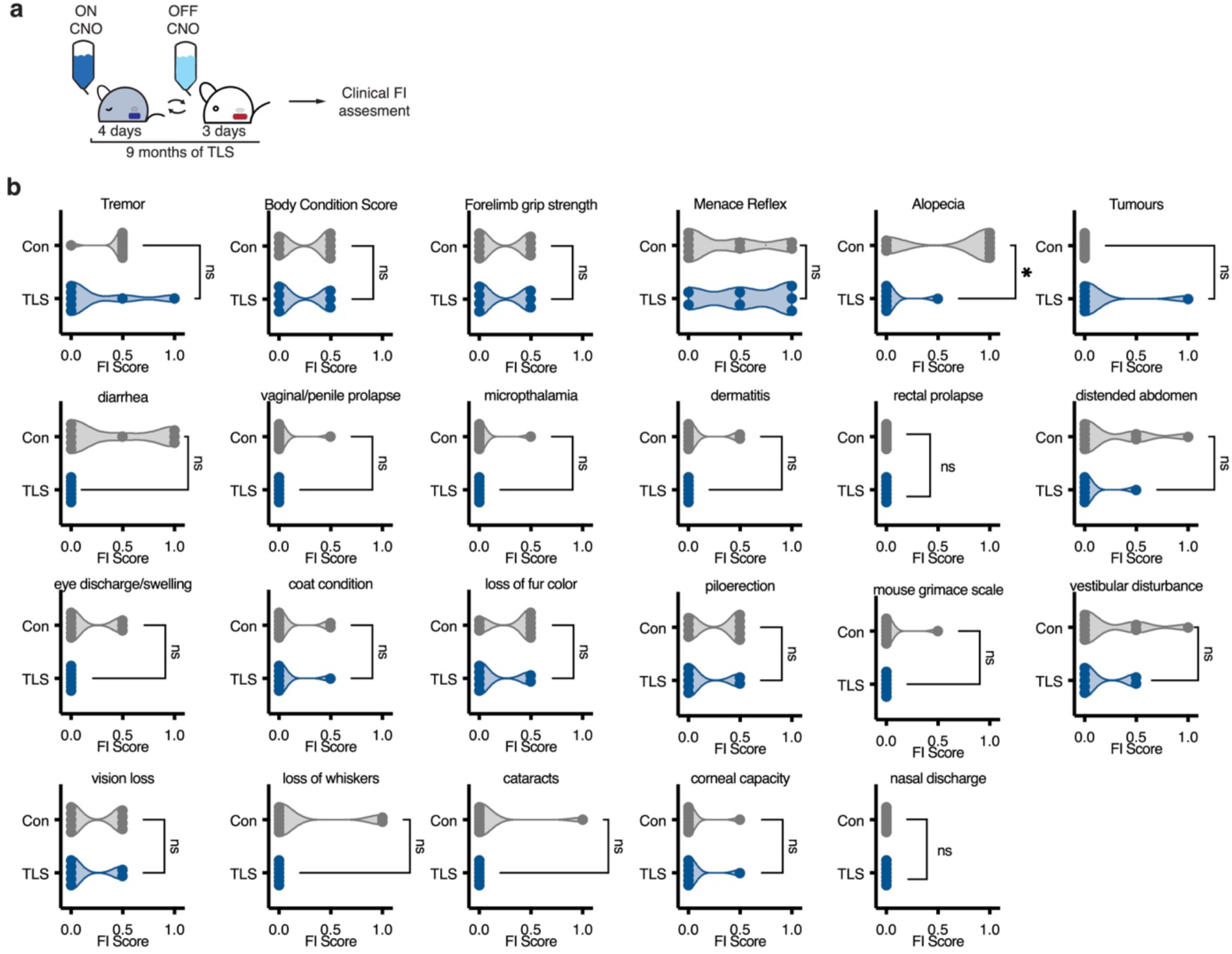
Clinical Frailty Index measurements. **a,** Schematic of TLS treatment and clinical FI assessment **b,** Frailty scores across the remaining 23 individual frailty index measurements of TLS and control mice after 9 months of TLS. TLS mice (0.07± 0.0) scored significantly lower than control mice (0.67± 0.0) on measurement of alopecia as determined by T-Test (\**P* = 0.01). Data reported as mean ± SEM.

**Extended Data Fig. 5.**
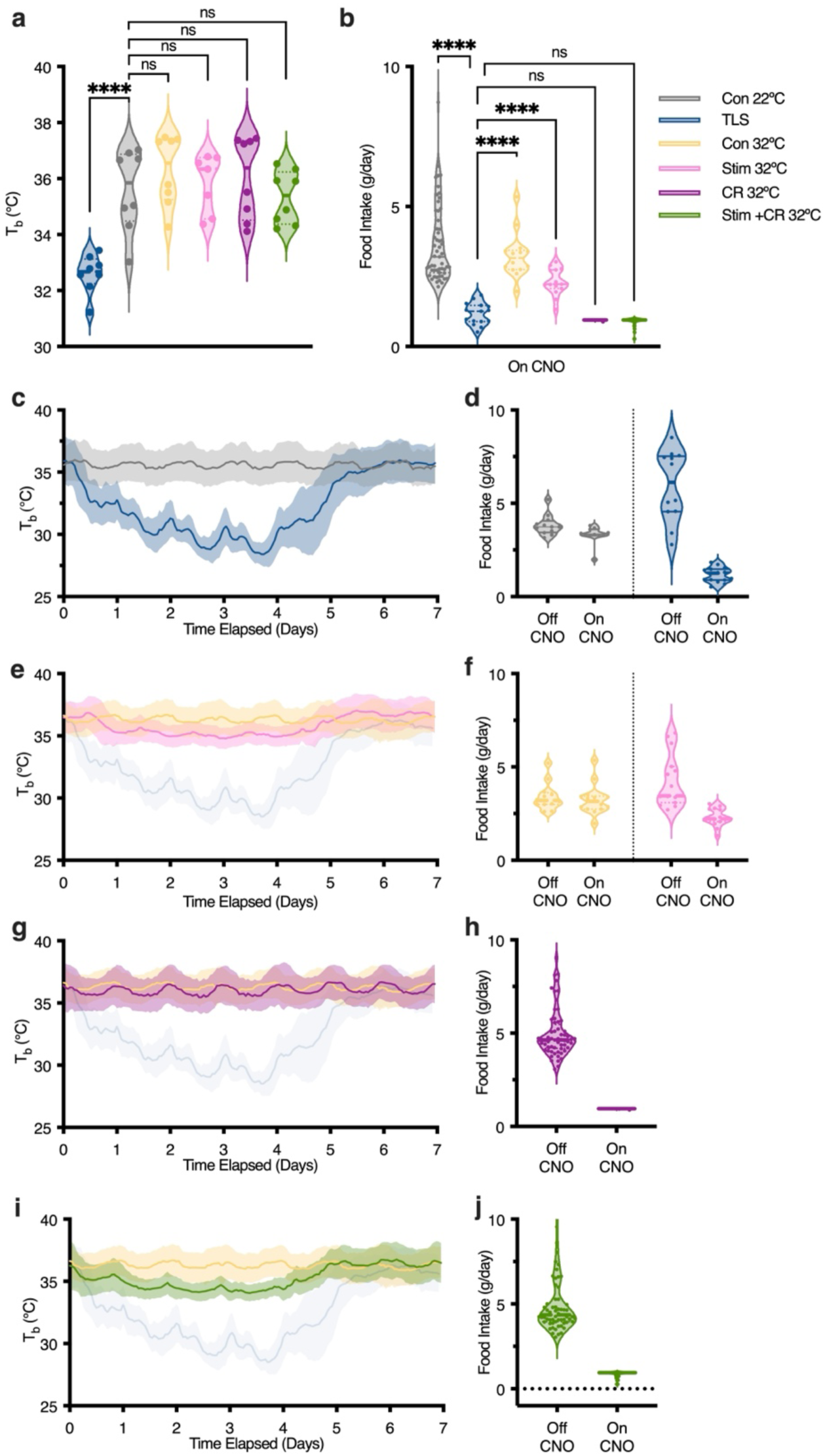
Food intake and T_b_ in TLS, *Con 22°C*, *Con 32°C, Stim 32°C*, *CR 32°C*, and *Stim + CR 32°C* mice. **a,** Average T_b_ of individual mice across all groups over 3 months. Only TLS mice had significantly lower T_b_ than Con 22°C mice (**** *P* < 0.0001). *Con 32°C* (ns, P = 0.649), *Stim 32°C* (ns, P = 0.994), *CR 32°C* (ns, *P* = 0.903), and *Stim+CR 32°C* (ns, *P* = 0.9915) mice all had similar average T_b_ to *Con 22°C* mice as determined by one-way ANOVA adjusted for multiple comparisons by Tukey’s HSD. **b,** Food intake across all groups for the duration of the experiment while on CNO. TLS mice (1.17±0.10 g) had significantly lower food intake than Con 22°C mice ( 3.67± 0.18 g) (****, *P* < 0.0001*), Con 32°C* mice (3.25±0.22 g) (****, *P* < 0.0001), and *Stim 32°C* mice (2.30± 0.12 g) (****, *P* < 0.0001). Importantly, TLS mice had similar food intake to *CR 32°C* mice (0.95± 0.0) (ns, *P* = 0.6274) and *Stim + CR 32°C* mice (0.88± 0.02) (ns, *P* = 0.413). **c,** Aggregate plot of T_b_ over 12-week experiment displayed over a 1-week of interval of control (grey) and TLS (blue) mice. Data plotted as mean ±SD. **d,** Food intake of TLS mice and Control mice while on and off CNO over duration of the experiment. **e,** Aggregate plot of T_b_ over 12-week experiment displayed over a 1-week interval of *Con 32°C* and *Stim 32°C* mice. TLS mice shown in blue for reference. Data plotted as mean ± SD. **f,** Food-intake of *Con 32°C* and *Stim 32°C* mice while on and off CNO. **g-h,** Characterization of *CR 32°C* mice, data shown as in **e-f**. **i-j,** Characterization of *Stim+CR 32°C* mice, data shown as in **e-f.**

**Extended Data Fig. 6.**
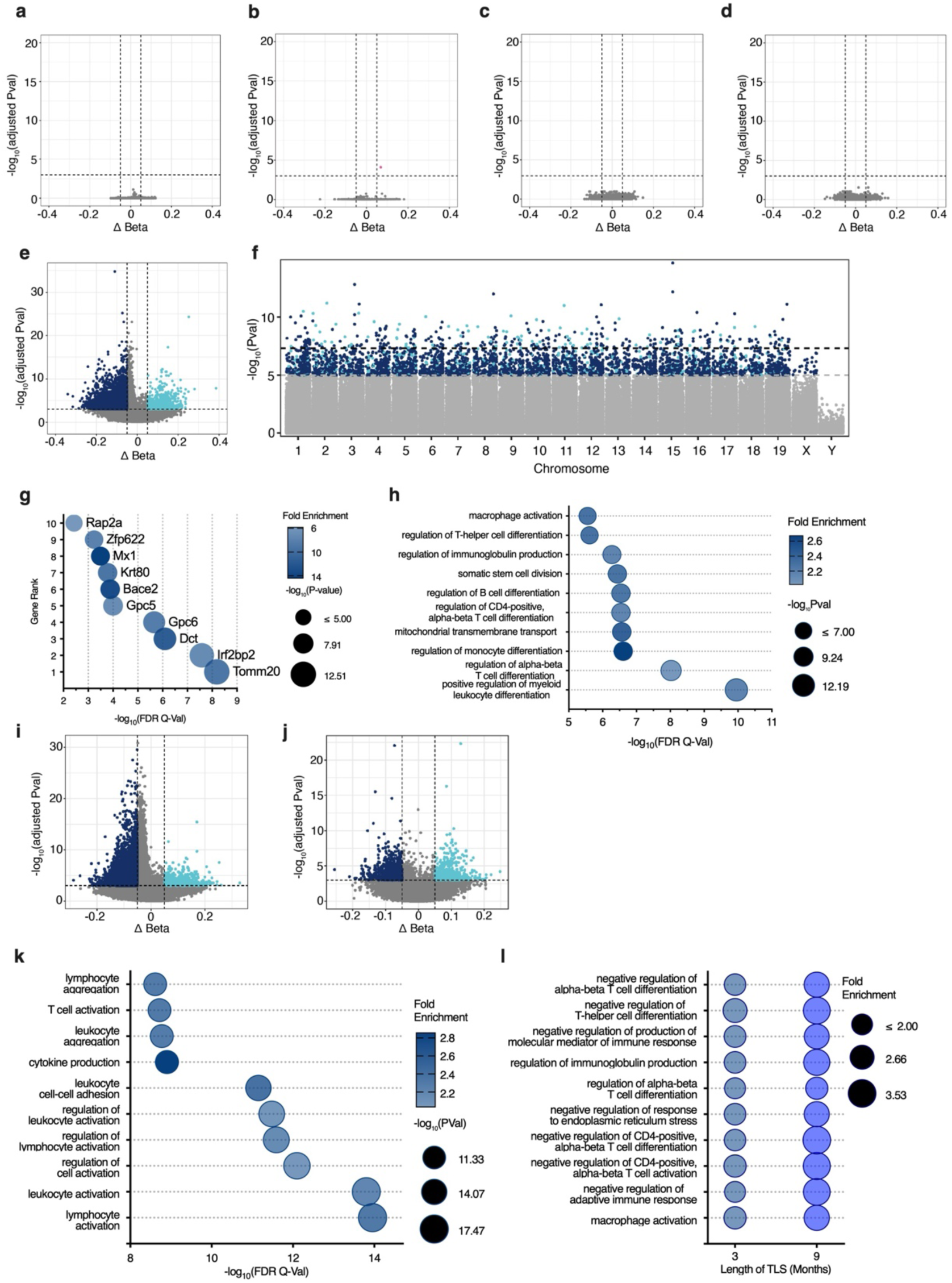
Differential methylation analysis. **a-e,** Volcano plots of differentially methylated regions across all groups as compared to Control 22°C mice. Differential methylation analysis was performed using the SeSAMe pipeline, which identified 183,635 genomic regions with correlated CpGs from 326,723 probes. Dashed lines represent significance thresholds (adjusted P-value < 10^-3^ and |ΔBeta| > 0.05). Importantly, only mice that underwent TLS had a meaningful number of differentially methylated regions as compared to Control 22°C mice after 3 months of treatment, suggesting temperature-dependent epigenetic remodeling. **a,** Control 32°C, no significantly differentially methylated regions **b,** Stim 32°C, 1 hypermethylated region (highlighted in pink). **c,** CR 32°C, no differentially methylated regions **d,** Stim+CR 32°C, had no differentially methylated regions **e,** TLS, 701 significantly hypermethylated regions highlighted in light blue, 5,332 significantly hypomethylated regions highlighted in dark blue. **f,** Manhattan plot displaying the 286,212 probes that mapped to the mouse genome using the SeSAME pipeline to visualize the genomic locations of differentially methylated probes between TLS and Control 22°C mice. Lower dashed line represents the significance threshold (raw P-value < 10^-5^). Probes that were significantly hypermethylated in TLS mice as compared to controls are highlighted in light blue; significantly hypomethylated probes are highlighted in dark blue. **g,** Top 10 enriched gene hits in TLS mice as compared to Control 22°C mice after 3 months of treatment identified via Genomic Regions Enrichment of Annotations Tool (GREAT) analysis from all differentially methylated regions. **h,** Top 10 enriched biological processes in TLS mice as compared to Control 22°C mice after 3 months of treatment identified via GREAT analysis. Only biological processes with more than 10 foreground gene hits were included. **i-j,** Volcano plots of differentially methylated regions in TLS mice as compared to control mice after 6 **(i)** and 9 months of TLS **(j)**. Differential methylation analysis was performed using the SeSAMe pipeline, which identified 183,635 correlated genomic segments. Dashed lines represent significance thresholds (adjusted P-value < 10^-3^ and |ΔBeta| > 0.05). Significantly hypermethylated regions are highlighted in light blue; significantly hypomethylated regions are highlighted in dark blue. **i,** After 6 months of TLS, there were 567 significantly differentially hypermethylated regions and 10,614 significantly differentially hypomethylated regions. **j,** After 9 months of TLS, there were 517 significantly differentially hypermethylated regions and 871 significantly differentially hypomethylated regions. **k,** Top 10 enriched biological processes after 9 months of TLS identified via GREAT analysis. Only biological processes with more than 10 foreground gene hits were included. **l,** Shared significantly enriched biological processes after 3 and 9 months of TLS. Biological processes were limited to those with an FDR < 0.05 and more than 5 foreground gene hits.

## Notes

### Competing Interest Statement

S. Horvath and R.T. Brooke are founders of the non-profit Epigenetic Clock Development Foun-dation, which licenses several patents from UC Regents including a patent on the mammalian methylation array platform. These patents list S.Horvath as inventor. V.G.S. serves as an advisor to and/or has equity in Branch Biosciences, Ensoma, and Cellarity, all unrelated to this work.

